# Helminth infection impacts hematopoiesis

**DOI:** 10.1101/2023.02.10.528073

**Authors:** Tobias Wijshake, Jipeng Wang, Joseph Rose, Madeleine Marlar-Pavey, James J. Collins, Michalis Agathocleous

## Abstract

Helminth infections are common in animals. However, the impact of a helminth infection on the function of hematopoietic stem cells (HSCs) and other hematopoietic cells has not been comprehensively defined. Here we describe the hematopoietic response to infection of mice with *Schistosoma mansoni,* a parasitic flatworm which causes schistosomiasis. We analyzed the frequency or number of hematopoietic cell types in the bone marrow, spleen, liver, thymus, and blood, and observed multiple hematopoietic changes caused by infection. Schistosome infection impaired bone marrow HSC function after serial transplantation. Functional HSCs were present in the infected liver. Infection blocked bone marrow erythropoiesis and augmented spleen erythropoiesis, observations consistent with the anemia and splenomegaly prevalent in schistosomiasis patients. This work defines the hematopoietic response to schistosomiasis, a debilitating disease afflicting more than 200 million people, and identifies impairments in HSC function and erythropoiesis.

## Introduction

Schistosomiasis is a parasitic disease caused by infection with *Schistosoma* flatworms. It afflicts more than 200 million people in Africa, the Middle East, South-East Asia, and South America (1, 2). Vaccines are not available. Treatment can clear adult parasites but is ineffective against immature parasites, does not prevent reinfection, nor does it reverse multi-organ immunopathology (3). As a result, the chronic symptoms of schistosomiasis contribute substantially to the global disability burden, creating a cycle of poverty and infection (4–7). Schistosomes are shed as larvae from *Biomphalaria* freshwater snails, infect humans by penetrating the skin, and can live in the circulation for decades (8), continuously laying eggs which lodge in liver, bladder, and other organs. Egg antigens trigger a Th2 response which dominates the chronic phase of the disease and is the main cause of pathology (1, 3, 9, 10). Several schistosomiasis symptoms including anemia, splenomegaly, and chronic inflammation suggest hematopoietic involvement. However, the effect of schistosomiasis on HSCs and restricted progenitors is mostly uncharacterized.

Until recently most humans were likely to be parasitized (11). The prevalence of infection with schistosomes or other helminths was as high as 50% in some populations before the modern era (12), or in modern untreated populations living in endemic areas (13). Primates and other animals are parasitized in the wild (14) and in some areas as many as 90% of primates have a history of schistosomiasis (15). Thus, HSCs and the hematopoietic system have likely evolved under near-constant pressure from schistosomes and other helminths. Recent studies have examined the impact on HSC function of various infections (16), including *Mycobacterium avium,* (17, 18)*, E. coli* (19), *Streptococcus* (20), *Plasmodium* (21), *Ehrlichia muris* (22), *Leishmania* (23), *C. albicans* (24), *Salmonella* (25, 26) and several viruses (27–29). Most of these infections induce a strong proinflammatory Th1 response. The impact of Th2 response-dominated chronic helminth infections on hematopoiesis has been much less characterized, (30, 31) and HSCs and progenitor responses in this context have not been systematically defined. To determine this, we examined the impact of schistosomiasis on HSC function and hematopoiesis.

## Results

### The effects of schistosome infection on hematopoiesis

To understand the impact of schistosomiasis on HSCs and hematopoietic progenitors, we infected mice with the human pathogen *Schistosoma mansoni.* In this model, eggs deposited in the liver trigger granuloma formation and schistosomiasis pathology starting from ~ 5 weeks post-infection. We analyzed the hematopoietic system of mice 7 weeks post-infection as compared to uninfected mice (Figure 1A). Bone marrow cellularity did not significantly change after infection (Figure 1B). The frequencies of CD150^+^CD48^-^Lineage^-^Sca-1^+^Kit^+^ HSCs and CD150^-^CD48^-^Lineage^-^Sca-1^+^Kit^+^ multipotent progenitors (MPPs) were unchanged (Figure 1C-D). Infection increased the frequency of CD150^-^CD48^+^Lineage^-^Sca-1^+^Kit^+^ hematopoietic progenitor cells (HPC-1) (Figure 1E) and decreased the frequency of CD34^+^CD16/32^-^Lineage^-^Sca-1^-^Kit^+^ common myeloid progenitors (CMPs) and CD34^-^CD16/32^-^Lineage^-^Sca-1^-^Kit^+^ megakaryocyte-erythroid progenitors (MEPs) (Figure 1F-G). The observed decline in CMP and MEP frequency was similar to a previous study examining the effects of schistosome infection in *Apoe*-deficient mice on a high-fat diet (32). The frequencies of CD150^+^CD48^+^Lineage^-^Sca-1^+^Kit^+^ (HPC-2) and CD34^+^CD16/32^+^Lineage^-^Sca-1^-^Kit^+^ granulocyte-monocyte progenitors (GMPs) did not change (Figure 1H-I). The spleen size and cellularity significantly increased after infection (supplemental Figure 1A-B), however the frequency of HSCs and most progenitor cell types in the spleen did not change (supplemental Figure 1C-P). This suggests that in contrast to other infectious or inflammatory challenges, the spleen is not a reservoir for multilineage hematopoiesis in schistosomiasis despite its increased size.

**Figure 1.**
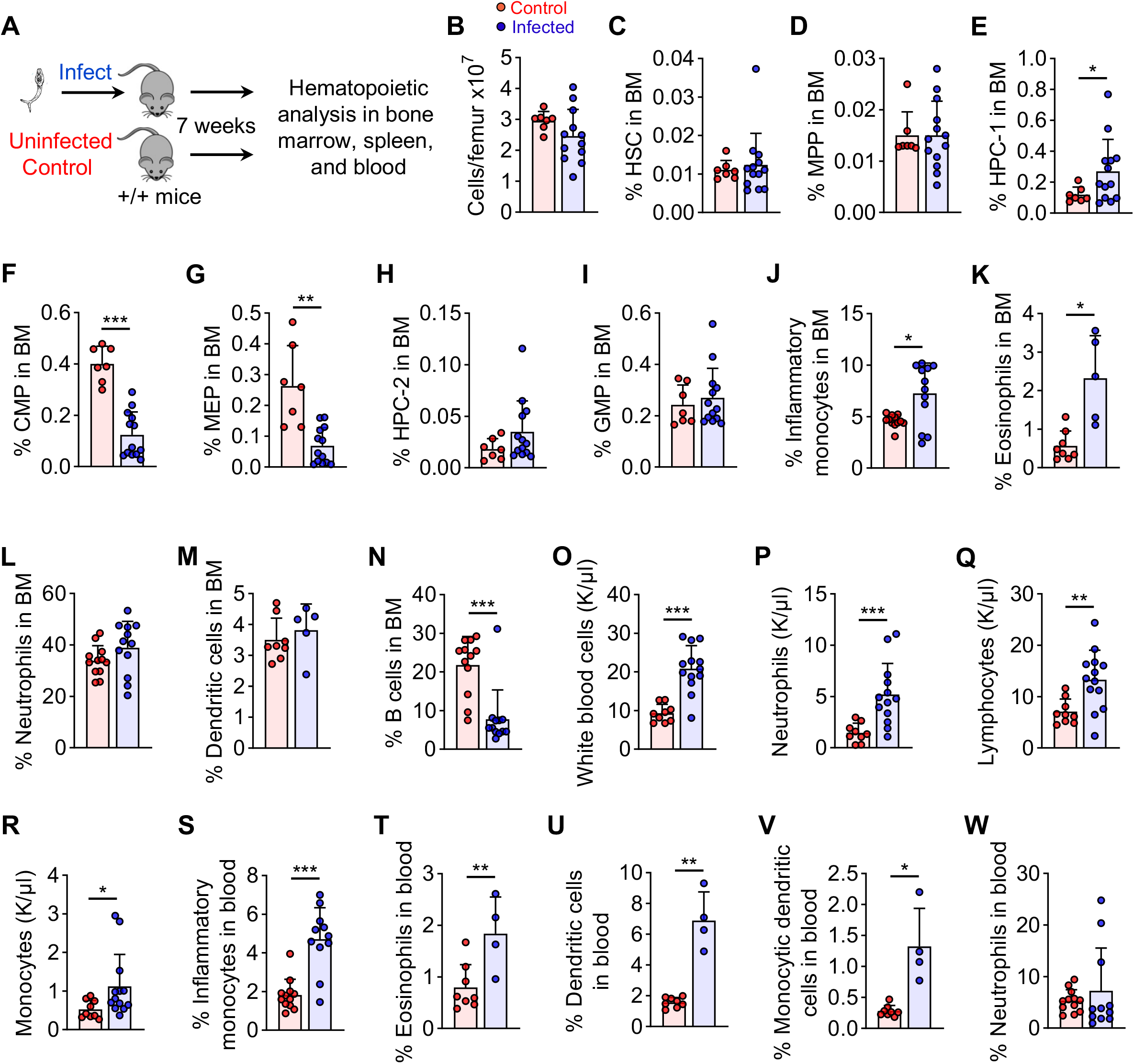
The effects of schistosome infection on the frequency of hematopoietic cells in the bone marrow and on blood cells. (A) Schematic overview of analysis of the hematopoietic and blood system 7 weeks after *Schistosoma mansoni* infection. (B-N) The frequency of HSCs, progenitors and mature cell types in the bone marrow of *S. mansoni* infected mice or uninfected controls (n = 7-13 mice per treatment). (O-R) Blood cell counts of white blood cells, neutrophils, lymphocytes, and monocytes of infected or uninfected mice (n = 9-13 mice per treatment). (S-W) The frequency of immune cell types in the blood of infected or uninfected mice (n = 4-11 mice per treatment). All graphs show mean ± s.d. *p < 0.05, **p < 0.01, ***p < 0.001. Statistical significance was assessed with a t-test with Welch’s correction (B-C, E, G-H, J-K, O-Q, S and U-W), a t-test (F, I, L-M, R and T), or a Mann-Whitney test (D and N).

In the myeloid lineage, infection preferentially increased the frequency of bone marrow monocytes and eosinophils but not neutrophils or dendritic cells (Figure 1J-M). Infection significantly decreased bone marrow B cell frequency (Figure 1N) in agreement with a recent report (33). The infected spleen had an increased frequency and number of most myeloid lineage cell types (supplemental Figure 1Q-AB). Infected mice had a higher white blood cell count than uninfected mice, and increased numbers or frequencies of neutrophils, monocytes, eosinophils, and dendritic cells in the blood (Figure 1O-W). The development of major T cell progenitor cell types in the thymus was not significantly impacted by infection (supplemental Figure 2). This contrasts with the severe impact of many other infections on thymus cellularity (34). To determine the effects of infection on T cells, we analyzed T-cell subpopulations in the blood, bone marrow, spleen, and liver. In the marrow, the frequency of total CD4^+^ and effector memory CD4^+^ T cells increased and the frequency of CD8^+^ T cells did not change (supplemental Figure 3A-F). Significant differences in T cell subsets between infected and uninfected mice were not observed in the spleen (supplemental Figure 3G-L). The frequency of naïve CD4^+^ T cells declined in the blood (supplemental Figure 3M-R). The liver had an increased frequency of total CD4^+^ or CD8^+^ T cells, particularly of a resident memory immunophenotype (supplemental Figure 3S-V). Therefore, schistosome infection caused several changes in the frequency of hematopoietic and immune cells in the marrow, spleen, blood, and liver.

### Infection impairs bone marrow HSC function after serial transplantation

To test if infection changes HSC function, donor bone marrow cells from infected or uninfected mice were mixed with competitor bone marrow cells from uninfected mice and transplanted into lethally irradiated recipients (Figure 2A). There was no significant difference in donor cell reconstitution capacity between infected and uninfected mice (Figure 2B-E; supplemental Figure 4A-B). At 16 weeks after transplant, there was no significant difference in reconstituted lineage composition between blood cells from infected as compared to uninfected mice (supplemental Figure 4C-G) suggesting infection did not cause long-term cell-intrinsic changes in HSC differentiation potential. Despite the maintenance of hematopoietic reconstitution in the peripheral blood, the bone marrow of transplant recipients had proportionately fewer infected donor-derived HSCs and some restricted progenitors as compared to uninfected donor-derived HSCs or progenitors (Figure 2F-G). This suggested an impairment in HSC self-renewal after transplantation. To test that, bone marrow cells from primary transplant recipients were transplanted into lethally irradiated secondary transplant recipients (Figure 2H). The blood reconstitution capacity of bone marrow cells from infected mice significantly decreased as compared to uninfected mice (Figure 2I-L; supplemental Figure 4H-I). The proportion of infected donor-derived myeloid progenitors and total hematopoietic cells in the marrow of secondary transplant recipients decreased (Figure 2M-N). Therefore, schistosome infection impaired HSC function after serial transplant.

**Figure 2.**
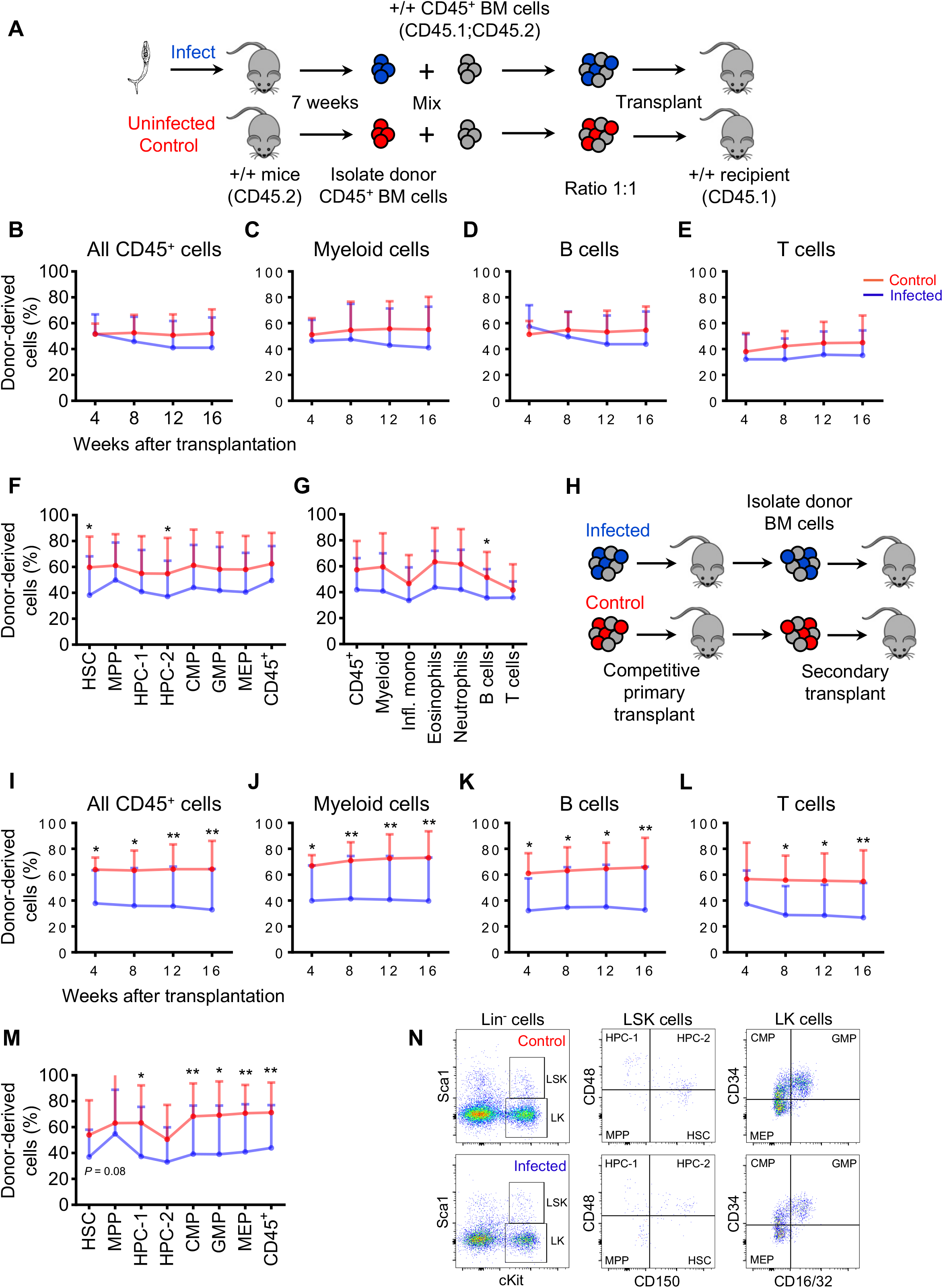
Schistosome infection impairs bone marrow HSC function. (A) Schematic overview of experiments to assess bone marrow HSC function after infection. 5 x 10^5^ CD45.2^+^ donor cells from bone marrow of mice infected with *S. mansoni* for seven weeks, or from uninfected mice, were mixed with 5 x 10^5^ CD45.1^+^CD45.2^+^ competitor bone marrow cells from uninfected mice and transplanted to each lethally irradiated CD45.1^+^ recipient mouse (n = 5 donor and 21-25 recipient mice per treatment). (B-E) Donor cell reconstitution of CD45^+^, myeloid, B, and T cells in the blood at the indicated time points. (F) The percentage of donor-derived hematopoietic stem and progenitor cells in the bone marrow (n = 21-25 mice per treatment). (G) The percentage of donor-derived myeloid, B and T cells in the bone marrow (n = 15 mice per treatment). (H) Schematic overview of the secondary transplantation. 1 x 10^7^ bone marrow cells from primary recipient mice were transplanted into each lethally irradiated secondary recipient mouse (n = 3 donor mice and 14-15 recipient mice per treatment). (I-L) Donor cell reconstitution of CD45^+^, myeloid, B and T cells in the blood at the indicated time points after secondary transplantation. (M) The percentage of donor-derived hematopoietic stem and progenitor cells in the bone marrow after secondary transplantation (n = 13-14 mice per treatment). (N) Representative flow plots of donor-derived Lineage^-^ cells (left), Lin^-^Sca-1^+^Kit^+^ (LSK) cells (middle), and Lin^-^Sca-1^-^Kit^+^ (LK) cells (right). All graphs show mean ± s.d. *p < 0.05, **p < 0.01, ***p < 0.001. Statistical significance was assessed with a repeated measures mixed model (B-E, and I-L), a t-test (F-G, and M, HSC and HPC-2 cells), a t-test with Welch’s correction (M, HPC-1; CMP; GMP; MEP; and CD45^+^ cells) or a Mann-Whitney test (F, HPC-1 cells; and M, MPP cells).

### Hematopoietic activity in the liver of infected mice

The liver of infected mice harbors schistosome eggs which trigger granuloma formation. To test if schistosomiasis elicits multilineage hematopoiesis in the liver, we analyzed the frequency of phenotypic HSCs and other hematopoietic progenitors 7 weeks after infection with *S. mansoni.* The liver of infected mice had an increased frequency of phenotypic HSCs, HPC-1, HPC-2, CMPs, GMPs and MEPs (Figure 3A-F). Among mature cells, the frequency of inflammatory monocytes, eosinophils, dendritic cells, and monocytic dendritic cells also increased after infection, the frequency of neutrophils was unchanged, and the frequency of B cells decreased (Figure 3G-L). The infected liver was enlarged (Figure 3M). To test if the increased frequency of phenotypic HSCs corresponded to an increase in HSC function, we transplanted 2 x 10^6^ cells from the liver of infected or uninfected donor mice in competition with 4 x 10^5^ bone marrow cells from uninfected mice into lethally irradiated recipient mice (Figure 3N). The peripheral blood of recipient mice contained significantly more donor-derived cells from infected livers than from uninfected livers (Figure 3O-R). Nine out of ten mice receiving donor cells from infected liver showed donor-derived multilineage reconstitution as compared to only one out of fourteen mice receiving donor cells from uninfected liver (Figure 3S). The bone marrow of transplant recipients had a significantly higher proportion of HSCs and restricted progenitors derived from infected donor livers as compared to uninfected donor livers (Figure 3T-U). Thus, functional HSCs were present in the liver of infected mice.

**Figure 3.**
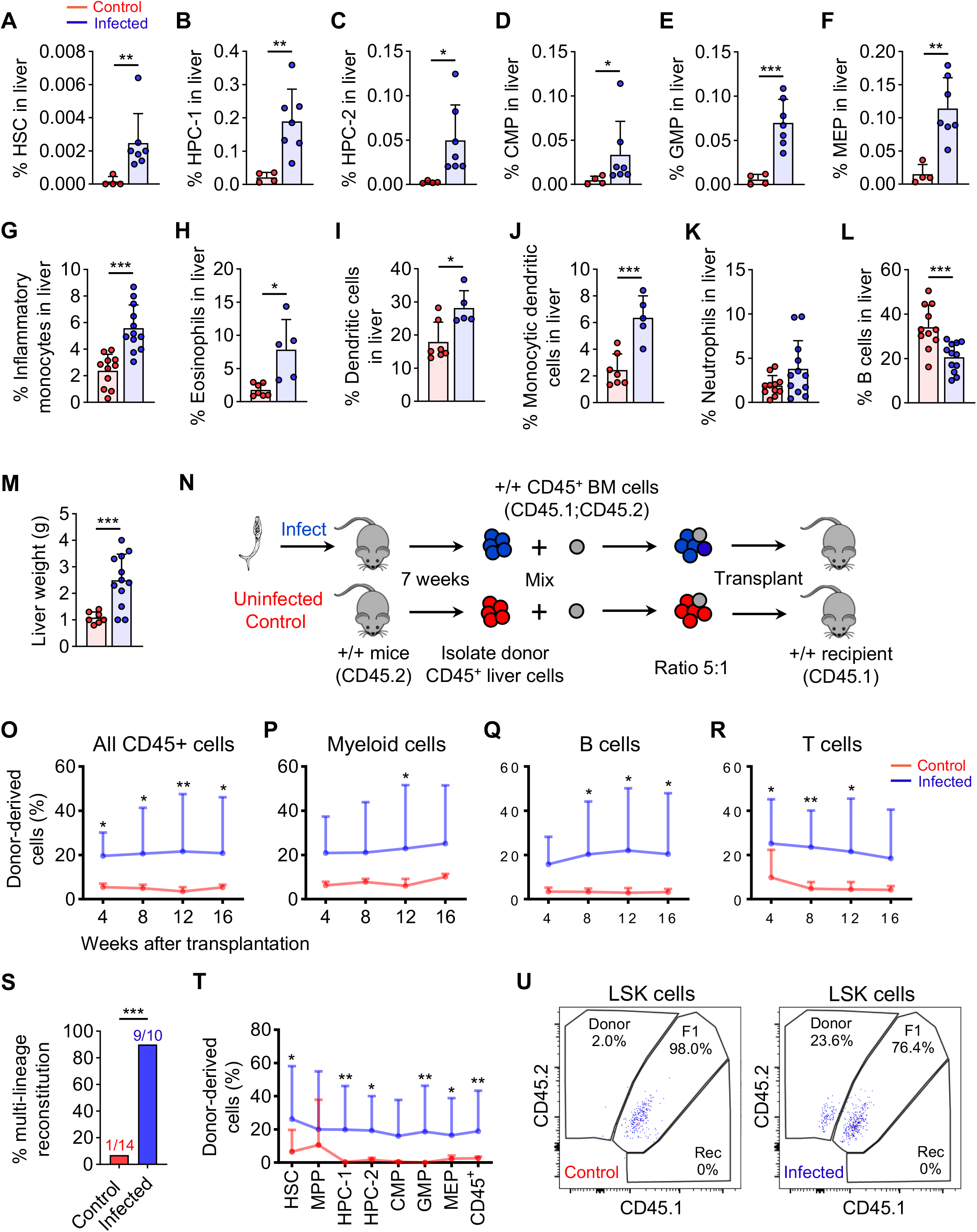
Schistosome infection stimulates liver hematopoiesis. (A-L) Frequency of hematopoietic stem, progenitor and mature immune cells in the liver of schistosome infected or uninfected control mice (n = 4-12 mice per treatment). (M) Liver weight after schistosome infection. (N) Schematic overview of experiments to assess HSC function in the infected liver. 2 x 10^6^ CD45^+^ donor cells from livers of infected or uninfected control mice were mixed with 4 x 10^5^ CD45^+^ competitor bone marrow cells from uninfected mice and transplanted to each lethally irradiated recipient mouse (n = 3 donor and 10-14 recipient mice per treatment). (O-R) Donor cell reconstitution of CD45^+^, myeloid, B and T cells in the blood at the indicated time points after transplantation. (S) The proportion of recipient mice which showed multilineage reconstitution after competitive transplantation of donor CD45^+^ cells from the liver of infected mice or uninfected controls. Multilineage reconstitution was defined as > 2% donor cell chimerism in peripheral blood myeloid, B and T cells at 16 weeks after transplantation. (T) The percentage of donor-derived hematopoietic stem and progenitor cells in the bone marrow after transplantation (n = 10-14 mice per treatment). (U) Representative flow plots of chimerism analysis of LSK cells. All graphs show mean ± s.d. *p < 0.05, **p < 0.01, ***p < 0.001. Statistical significance was assessed with a Mann-Whitney test (A, T, HSC; MPP; HPC-1; and GMP), a t-test with Welch’s correction (B-C, E, H, K, M, and T, HPC-2; CMP; and CD45^+^), a t-test (D, F-G, I-J, L, and T, MEP), a repeated measures mixed model (O-R), and a Fisher’s exact test (S).

### Schistosome infection blocks marrow erythropoiesis and increases spleen erythropoiesis

Schistosome-infected mice were anemic and thrombocytopenic (Figure 4A-D). They also showed increased red blood cell distribution width, a marker of anemia and inflammation which in humans correlates with increased mortality (35) (Figure 4E). Chronic anemia is one of the most prevalent and disabling symptoms of schistosomiasis (36). Several causes have been proposed including anemia of inflammation, blood loss, erythrocyte spleen sequestration, or autoimmunity (36). Schistosome-infected mice showed a sharp decrease in CD71^+^Ter119^+^ erythroid progenitors and an accumulation of CD71^mid^Ter119^-^ immature progenitors in the marrow (Figure 4F-I). This is consistent with a block in marrow erythropoiesis. In contrast the spleen had an increased frequency and number of CD71^+^Ter119^-^ and CD71^+^Ter119^+^ erythroid progenitors (Figure 4J-M). Therefore, erythropoiesis in schistosomiasis infection shifted from the bone marrow to the spleen.

**Figure 4.**
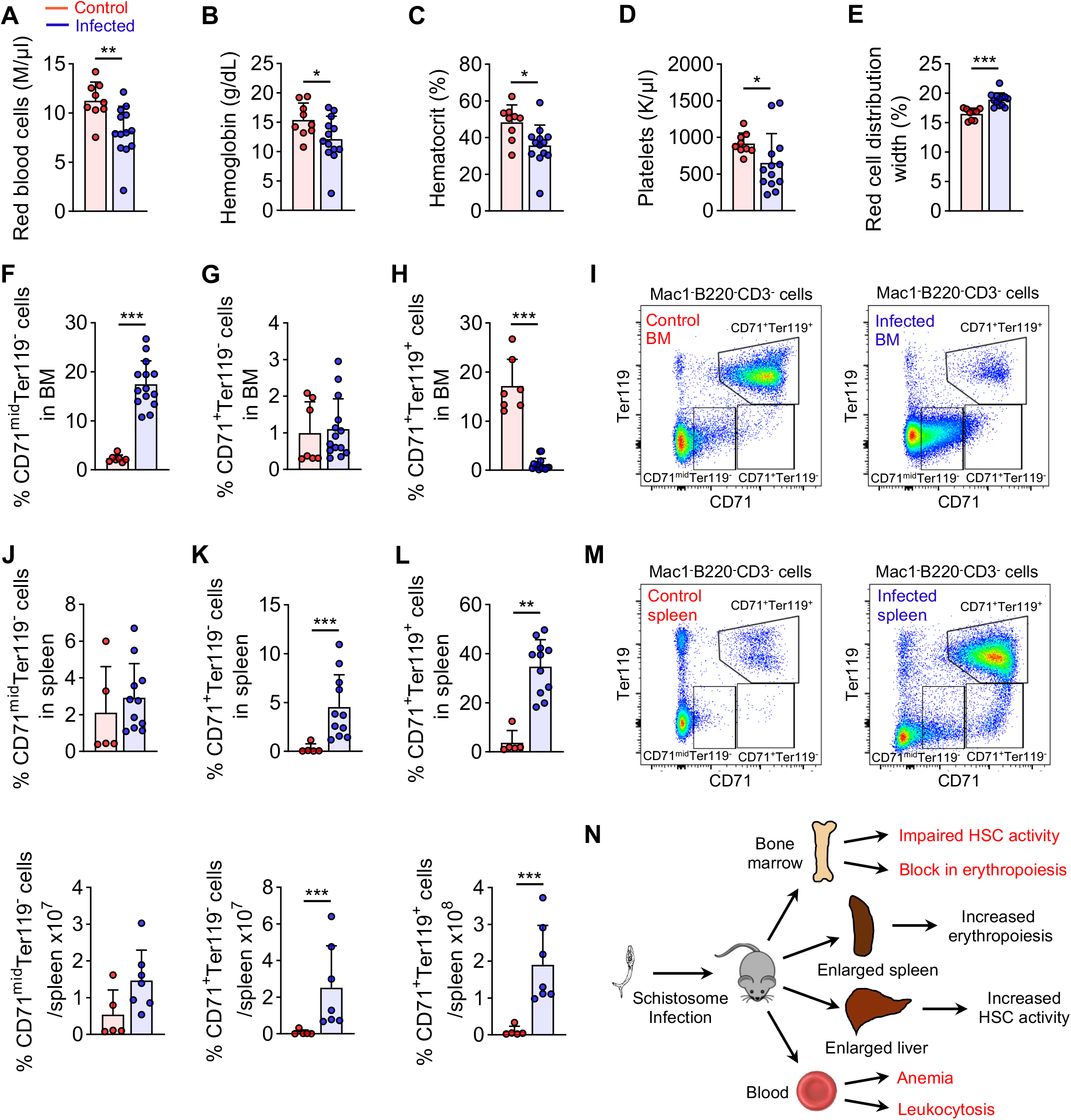
Schistosome infection blocks bone marrow erythropoiesis and increases spleen erythropoiesis. (A-E) Analysis of red blood cells, hemoglobin, hematocrit, platelets, and red blood cell distribution width of *S. mansoni* infected or uninfected mice. Analysis was performed 7 weeks after infection. (F-H) The frequency of CD71^mid^Ter119^-^ immature progenitors, CD71^+^Ter119^-^ and CD71^+^Ter119^+^ erythroid progenitors in the bone marrow of infected mice or uninfected control mice (n = 7-14 mice per treatment). (I) Representative flow plots of progenitors quantified in (F-H). (J-L) The frequency of CD71^mid^Ter119^-^ immature progenitors, CD71^+^Ter119^-^ and CD71^+^Ter119^+^ erythroid progenitors in the spleen of infected mice or uninfected control mice (n = 5-11 mice per treatment). (M) Representative flow plots of progenitors quantified in (J-L). (N) Graphical summary. All data show mean ± s.d. *p < 0.05, **p < 0.01, ***p < 0.001. Statistical significance was assessed with a t-test (A-C, E, G, and J-K) and a t-test with Welch’s correction (D, F, H, and L).

## Discussion

### The impact of schistosomiasis on hematopoietic stem cells

Our findings provide a framework to understand how *S. mansoni* infection affects hematopoiesis (Figure 4N). The frequency of bone marrow HSCs is not reduced after infection, and HSCs from infected mice can reconstitute primary recipients but are compromised in their ability to reconstitute secondary recipients. This suggests that schistosomiasis does not acutely impair HSC function but impairs the ability of the hematopoietic system to regenerate after repeated challenge. Populations in areas endemic for schistosomiasis have a high burden of anemia and of other infectious diseases, including malaria. This suggests that a reduction in HSC function after a schistosome infection could contribute to a long-term reduced capacity for hematopoietic regeneration after repeated infectious challenge.

### Erythropoiesis is blocked in the marrow but not the spleen

It is thought that inflammation is the most common cause of anemia in schistosomiasis (37). Our results suggest that anemia in schistosomiasis is partly caused by a sharp block in bone marrow erythropoiesis. This block is accompanied by a striking increase in spleen erythropoiesis. Inflammation is known to reduce marrow erythropoiesis (38) and consistent with this schistosomiasis increased inflammatory monocytes in the bone marrow and blood (Figure 1J, S). It is interesting that schistosomiasis arrested erythropoiesis in the marrow but not in the spleen. Our results suggest that an important function of the spleen is to make erythrocytes in schistosome infections. Because the infected spleen does not harbor more HSCs or myeloid progenitors than the uninfected spleen, splenomegaly in schistosomiasis is not a consequence of general extramedullary hematopoiesis but specifically of splenic erythropoiesis. In severe schistosomiasis, splenectomy is often used to alleviate hepatosplenomegaly and ensuing portal hypertension (39). Splenectomy increases risks for infections from bacterial or other parasitic infections (40, 41). It will be interesting to test if ameliorating the block in bone marrow erythropoiesis reduces splenomegaly and prevents anemia.

### Hematopoiesis in the liver

We show that HSCs and restricted progenitors are present in the liver after infection, as assayed by immunophenotypic and transplantation experiments. CD45^+^ cells from liver of infected mice competed against bone marrow cells from uninfected mice at a 5:1 donor:competitor ratio produced an average reconstitution of 20%, or 1:4 donor:competitor cells after transplant (Figure 3O). This suggests that functional HSC frequency is 20-fold lower in infected liver than bone marrow. Given that the bone marrow typically contains more CD45^+^ cells than the liver, systemic output of blood cells from liver HSCs in infection is likely to be negligible as compared to bone marrow hematopoiesis. However local production of immune cells in liver may be important for the immune response to granuloma formation. The infected liver harbors early B cell progenitors and restricted myeloid progenitors as assayed by *in vitro* and spleen colony-forming assays (42–47). Proliferating myeloid lineage cells are found in the periphery of granulomas (48). The production of monocyte-derived macrophages in the infected liver has been suggested to be protective (49–51). The fact that HSC or immature progenitor frequency is not elevated in spleen of infected mice suggests some specificity to liver hematopoiesis as opposed to general activation of extramedullary hematopoiesis. Are HSCs in the infected liver supported by specialized niches? Hepatic stellate cells can express known HSC niche factors, including SCF after schistosome infection (52) or CXCL12 in other contexts (53). Hepatic stellate cells serve as the major HSC niche cell type in fetal liver hematopoiesis by secreting SCF (54). Thus, we hypothesize that during schistosome infection HSCs may be supported by reactivation of a dormant liver hematopoietic niche.

## Methods

### Mice

Mice were on a C57BL/Ka background. Both male and female mice were used in all studies. Young adult mice were infected at the ages of 10-21 weeks and were either analyzed or used as donors for transplantation at 7 weeks after schistosome infection. C57BL/Ka-Thy-1.1 (CD45.2) and C57BL/Ka-Thy-1.2 (CD45.1) mice were used for transplantation experiments.

Mice were housed in the Animal Resource Center of UT Southwestern and all procedures were approved by the UT Southwestern Institutional Animal Care and Use committee.

### Schistosome infection

Each mouse was infected with around 200 *Schistosoma mansoni* (NMRI strain) cercariae released from infected *Biomphalaria glabrata* snails (Schistosome Resource Center) by percutaneous tail exposure(55).

### Cell isolation and hematopoietic analysis

Bone marrow cells were obtained by flushing femurs and tibia with 25G needle, or crushing femurs, tibias, vertebrae, and pelvic bones with a mortar and pestle in staining medium consisting of Ca^2+^/Mg^2+^-free Hank’s balanced salt solution (HBSS; Gibco), supplemented with 2% heat-inactivated bovine serum (Gibco). Spleens and thymuses were mechanically dissociated by trituration in staining medium. Livers were enzymatically digested for 30 minutes at 37°C in 1.5 ml RPMI-1640 (Sigma), containing 250 μg/ml liberase (Roche) and 100 μg/ml DNase I (Roche). Cell suspensions were filtered through a 40 μm strainer. Cell number was assessed with a Vi-CELL cell viability analyzer (Beckman Coulter). Blood was collected by cardiac puncture using a 25G needle and mixed in a tube containing 5 μl 0.5M EDTA. Complete blood cell counts were determined using a hemavet HV950 (Drew Scientific). For hematopoietic analysis, 40 μl blood was lysed in 1 ml of ammonium chloride buffer (ACK; 155mM NH4Cl; 10 mM KHCO_3_; 0.1 mM EDTA) for 10 minutes at 4°C. Cells were incubated with fluorescently conjugated antibodies for 90 minutes on ice when using CD34 antibody or for 30 minutes at 4°C. Cells were washed with staining media and resuspended in staining media containing 1 μg/ml DAPI or 1 μg/ml propidium iodide for live/dead discrimination. Cell populations were defined with the following markers: CD150^+^CD48^-^Lineage^-^Sca-1^+^Kit^+^ hematopoietic stem cells (HSCs), CD150^-^CD48^-^Lineage^-^Sca-1^+^Kit^+^ multipotent progenitor cells (MPPs), CD150^-^CD48^+^Lineage^-^Sca-1^+^Kit^+^ hematopoietic progenitor cells (HPC-1), CD150^+^CD48^+^Lineage^-^Sca-1^+^Kit^+^ hematopoietic progenitor cells (HPC-2), CD34^+^CD16/32^-^Lineage^-^Sca-1^-^Kit^+^ common myeloid progenitors (CMPs), CD34^+^CD16/32^+^Lineage^-^Sca-1^-^Kit^+^ granulocyte–monocyte progenitors (GMPs), CD34^-^CD16/32^-^ Lineage^-^Sca-1^-^Kit^+^ megakaryocyte–erythroid progenitors (MEPs), Mac-1^+^ myeloid cells, Mac1^+^CD115^+^Ly6C^+^Ly6G^-^ inflammatory monocytes, Mac1^+^CD115^-^Ly6C^mid/high^SiglecF^+^ eosinophils, Mac1^+^CD115^-^Ly6C^mid^Ly6G^+^ neutrophils, CD11c^+^ dendritic cells (DCs), CD11c^+^Mac1^+^Ly6C^+^ monocytic DCs (moDCs), CD11c^+^Mac1^-^ Mac1^-^DCs, Mac1^-^B220^+^ B cells, Mac1^-^CD3^+^ T cells, Mac1^-^B220^-^CD3^-^CD71^mid^Ter119^-^ immature erythroid progenitors, Mac1^-^ B220^-^CD3^-^CD71^+^Ter119^-^ erythroid progenitors, and Mac1^-^B220^-^CD3^-^CD71^+^Ter119^+^ erythroid progenitors. The Lineage cocktail for HSCs and progenitors consisted of CD2, CD3, CD5, CD8, Ter119, B220, and Gr-1 antibodies. T cell progenitor populations in the thymus were defined with the following markers, after excluding Mac-1^+^, B220^+^, and Ter119^+^ cells: CD4^+^CD8^+^ double-positive (DP), CD3^+^CD4^+^CD8^-^ (CD4^+^ single positive, CD4^+^SP), CD3^+^CD4^-^CD8^+^ (CD8^+^SP), CD4^-^ CD8^-^ double-negative (DN), CD4^-^CD8^-^CD44^+^CD25^-^ (DN1), CD4^-^CD8^-^CD44^+^CD25^+^ (DN2), CD4^-^ CD8^-^CD44^-^CD25^+^ (DN3), CD4^-^CD8^-^CD44^-^CD25^-^ (DN4), and CD3^-^CD4^-^CD8^+^ immature single positive (ISP). Mature T cell populations were defined by the following markers, after excluding Mac-1^+^, B220^+^, and Ter119^+^ cells: CD4^+^ (CD4^+^ cells), CD4^+^CD44^-^CD62L^+^ (naïve CD4^+^ cells), CD4^+^CD44^+^CD62L^+^ (CD4^+^ central memory cells), CD4^+^CD44^+^CD62L^-^CD69^+^ (CD4^+^ resident memory cells), CD4^+^CD44^+^CD62L^-^CD69^-^CD103^-^ (CD4^+^ effector memory cells), CD8^+^ (CD8^+^ cells), CD8^+^CD44^-^CD62L^+^ (naïve CD8^+^ cells), CD8^+^CD44^+^CD62L^+^ (CD8^+^ central memory cells), CD8^+^CD44^+^CD62L^-^CD69^+^ (CD8^+^ resident memory cells), and CD4^+^CD44^+^CD62L^-^CD69^-^CD103^-^ (CD8^+^ effector memory cells). All antibodies used in experiments are listed in supplementary Table 1. Analysis and sorting were performed using the FACSAria flow cytometer (BD Biosciences) or a FACSCanto (BD Biosciences). Data were analyzed using FlowJo (Flowjo LLC) or FACSDiva (BD Biosciences).

### Bone marrow and liver reconstitution assays

Recipient mice (CD45.1) were irradiated using an XRAD 320 X-ray irradiator (Precision X-Ray) with two doses of 540 rad (1080 rad in total) delivered at least 3 hours apart. Bone marrow cells were injected into the retro-orbital venous sinus of anesthetized recipients. Seven weeks prior to transplant, donor mice were either infected with 200 cercariae or left uninfected (controls). For competitive transplants, 5 x 10^5^ CD45^+^-selected bone marrow cells from infected or from uninfected donor (CD45.2) mice and 5 x 10^5^ competitor (CD45.1;CD45.2) cells were mixed and transplanted by injection into the retro-orbital venous sinus of anesthetized recipients. Recipient mice were maintained on antibiotic water (Baytril 0.08 mg/ml) for 1 week pre-transplantation, and for 4 weeks after transplantation. Blood was obtained from the tail veins of recipient mice every four weeks for at least 16 weeks after transplantation. Red blood cells were lysed in ACK lysis buffer. The remaining cells were stained with antibodies against CD45.2, CD45.1, C11b (Mac1), CD115, Ly6G, Ly6C, CD45R (B220), and CD3 and analyzed by flow cytometry. For the secondary bone marrow transplants, 1 x 10^7^ bone marrow cells from primary recipients were transplanted into lethally irradiated CD45.1 secondary recipients. For the competitive liver reconstitution assays, 2 x 10^6^ CD45^+^-selected liver cells from infected or from uninfected donor (CD45.2) mice and 4 x 10^5^ bone marrow competitor (CD45.1;CD45.2) cells from uninfected mice were mixed and transplanted into lethally irradiated CD45.1 recipient mice. Bone marrow cells for analysis of transplant recipient mice or for secondary transplantations was obtained by crushing femurs, tibias, vertebrae, and pelvic bones.

### Statistical analysis

Most figure panels show the pooled results from mice we analyzed from multiple independent experiments. Mice were allocated to experiments randomly. For most experiments the operator was not blinded to the treatment. Uninfected littermate controls, or uninfected controls from litters of the same parental strains born a few days apart were used for experiments. Prior to analyzing the statistical significance of differences among treatments, we tested whether data were normally distributed and whether variance was similar among treatments. To test for normal distribution, we performed the Shapiro-Wilk test when 3 ≤ n < 20 or the D’Agostino & Pearson test when n ≥ 20. To test if variability significantly differed among treatments, we performed F-tests. If the data did not significantly (p < 0.01 for at least one treatment) deviate from normality, we used a parametric test, otherwise data were log-transformed and tested for a significant deviation from normality. If the log-transformed data passed normality, a parametric test was used on the transformed data. If both the untransformed and log-transformed data did not pass the normality test, a non-parametric test was used on the untransformed data. To assess the statistical significance of a difference between two treatments, we used a t-test for data that was normally distributed and had equal variability, or a t-test with Welch’s correction for data that was normally distributed and had unequal variability, or a Mann-Whitney test for data that was not normally distributed. To assess the statistical significance of differences between treatments when multiple measurements were taken across time, we used a repeated measures mixed-effects model for data for which some values were missing. To determine the statistical significance between treatments for the presence of multi-lineage reconstitution, we used a Fisher’s exact test.

## Acknowledgments

M.A. is a Cancer Prevention and Research Institute of Texas (CPRIT) scholar and an American Society of Hematology faculty scholar. This work was supported by CPRIT (RR180007), the American Society of Hematology Faculty Scholar award, and grants from the National Institutes of Health (R01DK125713) to M.A and from the Welch Foundation (I-1948-20180324) and the National Institutes of Health (R01AI121037, R01AI150776) to JJC. We thank the members of the Collins and the Agathocleous labs for discussions; Q. Ding for mouse colony management; and the Moody Foundation Flow Cytometry Facility for flow cytometry.

## Authorship

### Contribution

T.W. collected and analyzed data, and performed statistical analysis; J.P., T.R., M.M.P., and M.A. performed experiments and analyzed data; J.J.C. and M.A. designed research; M.A. wrote the manuscript with help of T.W.; and all authors read and approved the manuscript.

### Conflict-of-interest disclosure

The authors declare no competing interests.

Data are available on request from Michalis Agathocleous

---

**Supplemental Figure 1.**
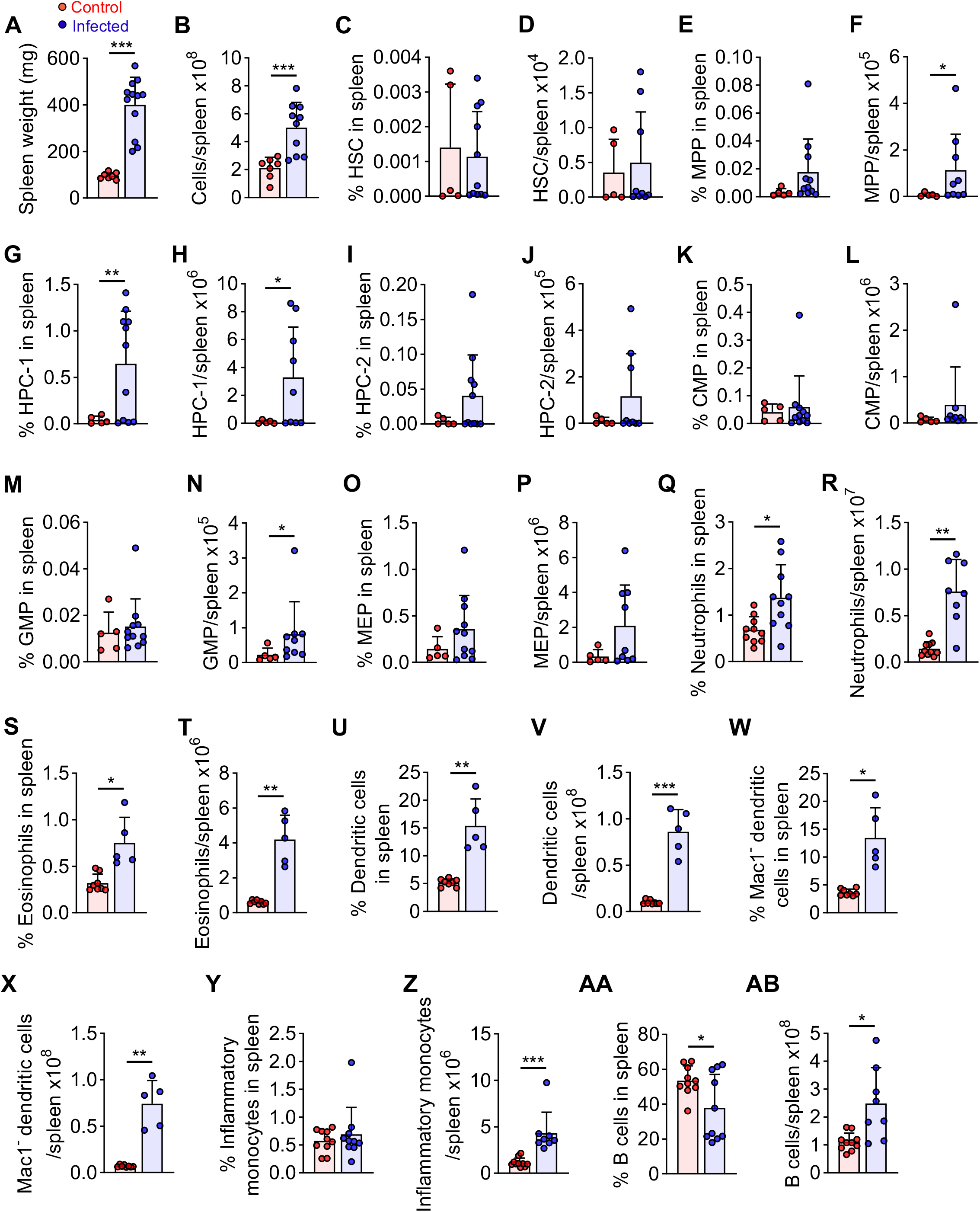
The effects of schistosome infection on hematopoietic and immune cells in the spleen. (A-B) Spleen weight and cell number after infection (n = 7-12 mice per treatment). (C-AB) The frequency of HSCs, progenitors and mature cell types in the spleen of infected mice or uninfected controls (n = 5-11 mice per treatment). All graphs show mean ± s.d. *p < 0.05, **p < 0.01, ***p < 0.001. Statistical significance was assessed with a t-test with Welch’s correction (A-B, G-H, P-U, W-X, and AA-AB), and a t-test (C-F, I-O, V, and Y-Z).

**Supplemental Figure 2.**
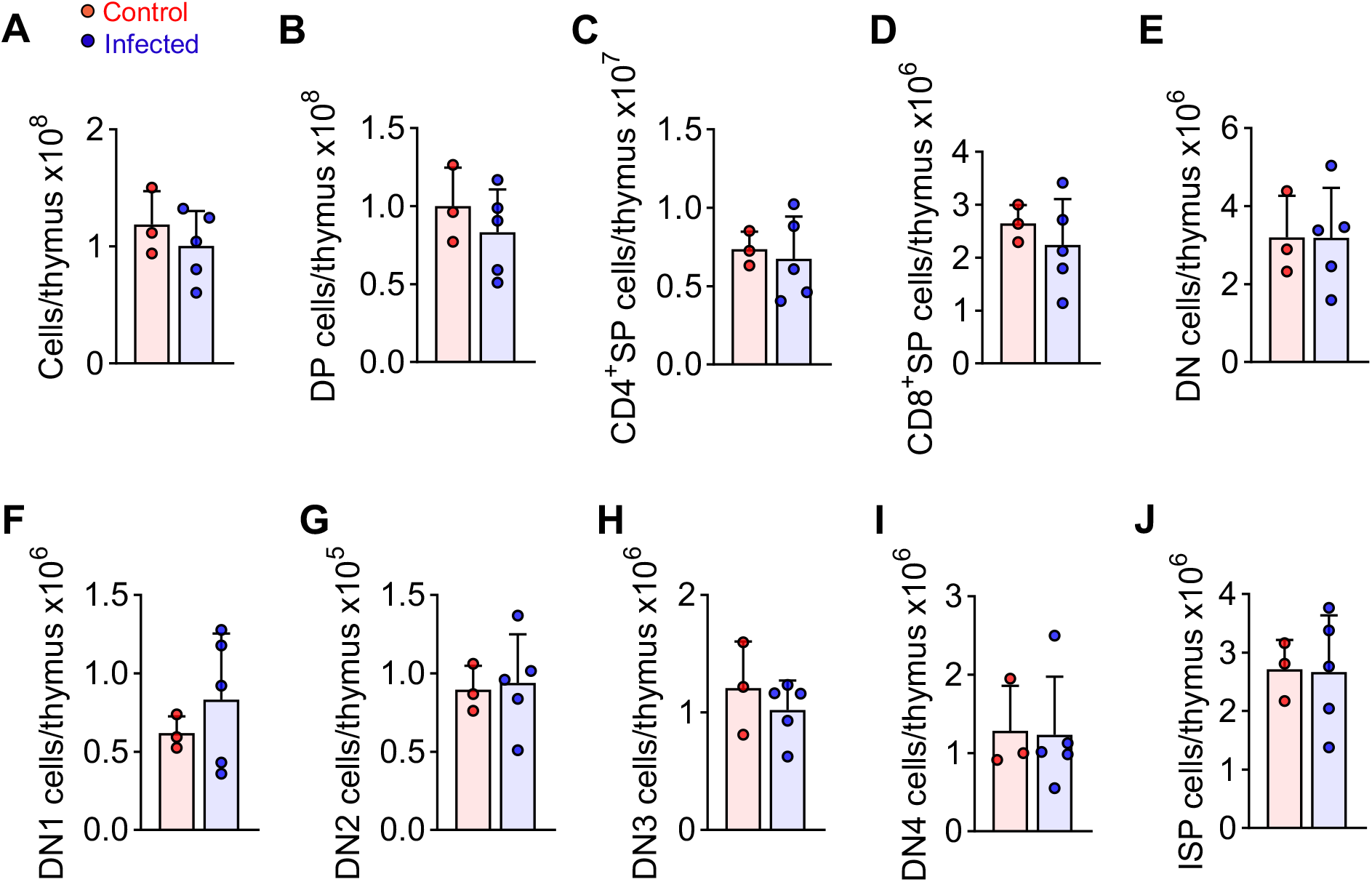
Schistosome infection does not affect the development of T cell progenitors in the thymus. (A) Thymus cellularity, and frequency of (B) CD4^+^CD8^+^ double-positive thymocytes, (C) CD4^+^ single-positive thymocytes, (D) CD8^+^ single-positive thymocytes, (E) CD4^-^CD8^-^ double-negative (DN) thymocytes, (F-I) subsets of double-negative thymocytes, and (J) CD3^-^CD8^+^ immature single-positive thymocytes in the thymus of infected mice or uninfected controls (n = 3-5 mice per treatment). All graphs show mean ± s.d.

**Supplemental Figure 3.**
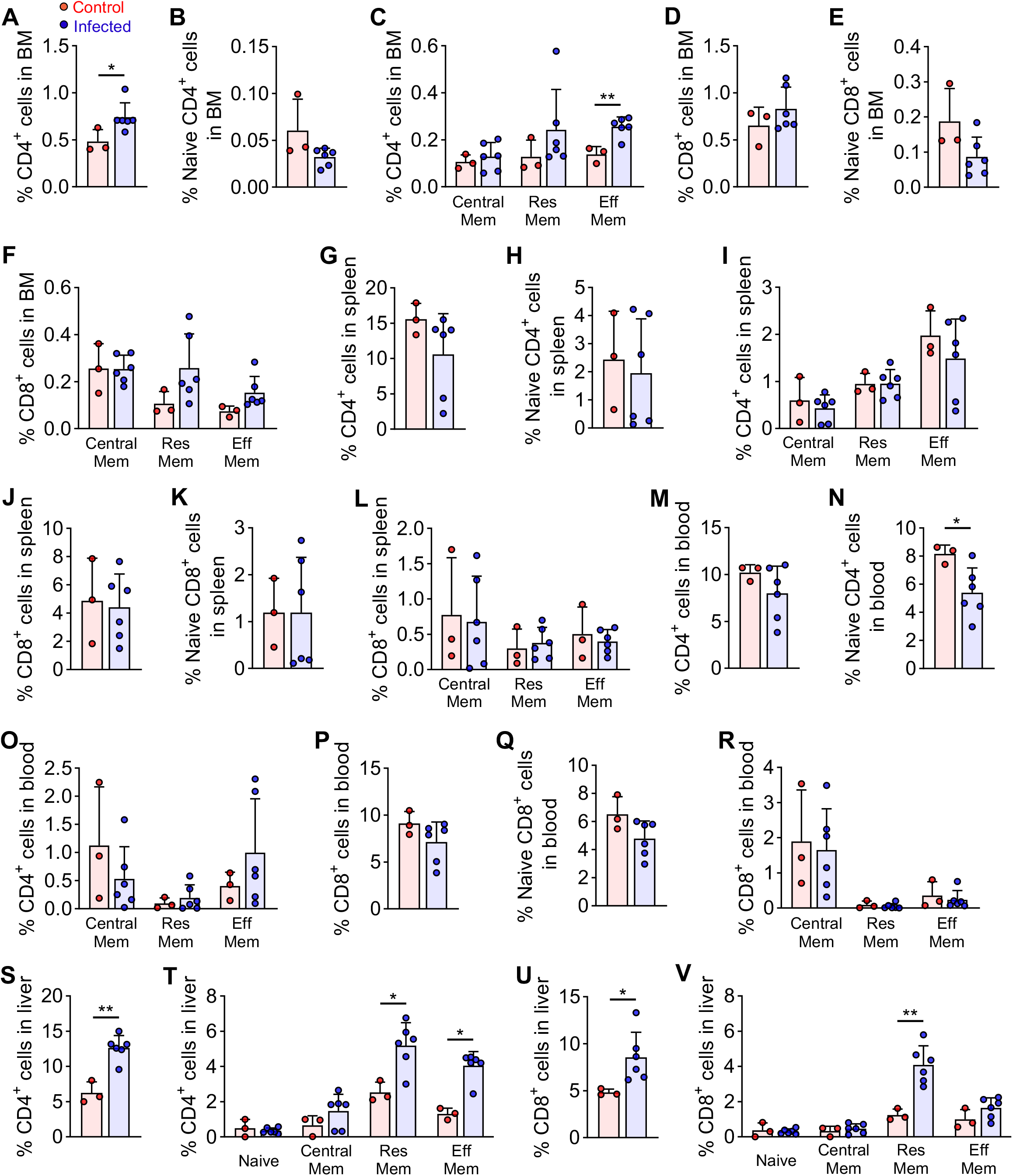
Effects of schistosome infection on the frequency of T cell subsets. (A-V) The frequency of CD4^+^ and CD8^+^ T cell subsets in the bone marrow, spleen, blood, and liver of infected mice or uninfected controls (n = 3-6 mice per treatment). All graphs show mean ± s.d. *p < 0.05, **p < 0.01. Statistical significance was assessed with a t-test (A, C, N, S, T, Res Mem; and V), a t-test with Welch’s correction (U), and a Mann-Whitney test (T, Eff Mem).

**Supplemental Figure 4.**
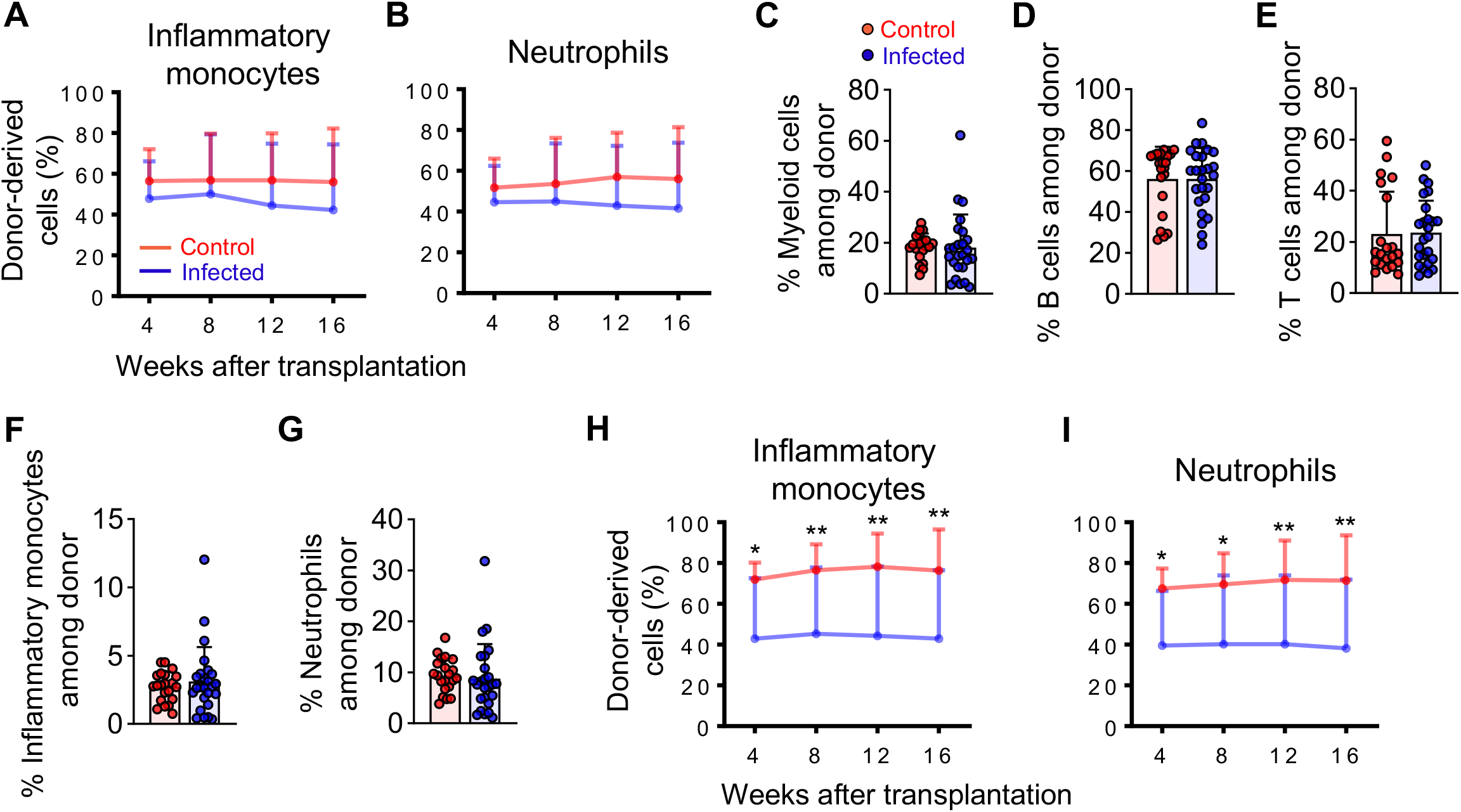
Effects of schistosome infection on bone marrow HSC function. (A-B) Donor cell reconstitution of inflammatory monocytes and neutrophils in the blood after competitive transplantation (n = 21-25 recipient mice per treatment). (C-G) Analysis of the frequency of immune cells among donor-derived CD45^+^ cells in the blood 16 weeks after transplantation (n = 21-25 mice per treatment). (H-I) Donor cell reconstitution of inflammatory monocytes and neutrophils in the blood after secondary transplantation (n = 14-15 recipient mice per treatment). All graphs show mean ± s.d. *p < 0.05, **p < 0.01. Statistical significance was assessed with a repeated measures mixed model (A-B and H-I), a t-test with Welch’s correction (C and F-G), and a t-test (D-E).

**Supplementary Table 1.** Antibodies.

| **Antibody** | **Company** | **Identifier** |
| --- | --- | --- |
| CD150 - PE | BioLegend | Cat#: 115904, clone TC15-12F12.2 |
| CD2 - FITC | Tonbo Biosciences | Cat#: 35-0021, clone RM2-5 |
| CD3 - FITC | BioLegend | Cat#: 100204, clone 17A2 |
| CD5 - FITC | BioLegend | Cat#: 100606, clone 53-7.3 |
| CD8a - FITC | Tonbo Biosciences | Cat#: 35-0081, clone 53-6.7 |
| Ly6G (Gr-1) - FITC | Tonbo Biosciences | Cat#: 35-5931, clone RB6-8C5 |
| Ter119 - FITC | Tonbo Biosciences | Cat#: 35-5921, clone TER-119 |
| CD45R (B220) - FITC | Tonbo Biosciences | Cat#: 35-0452, clone RA.3-6B2 |
| CD48 - AF700 | BioLegend | Cat#: 103426, clone HM48-1 |
| CD117 (c-Kit) - APC/Fire 750 | BioLegend | Cat#: 105838, clone 2B8 |
| Ly-6A/E (Sca-1) – PE/Cyanine7 | BioLegend | Cat#: 108114, clone D7 |
| CD34 – eFluor 660 | Invitrogen | Cat#: 50-0341-80, clone RAM34 |
| CD16/32 – BV510 | BioLegend | Cat#: 101333, clone 93 |
| CD115 (CSF-1R) – PE | BioLegend | Cat#: 135505, clone AFS98 |
| Ly-6G – FITC | BioLegend | Cat#: 127605, clone 1A8 |
| Ter119 – PE/Cy7 | BioLegend | Cat#: 116222, clone TER-119 |
| C11b (Mac1) – APC-Cyanine7 | Tonbo Biosciences | Cat#: 25-0112, clone M1/70 |
| Ly-6C – AF700 | BioLegend | Cat#: 128024, clone HK1.4 |
| CD71 – APC | BioLegend | Cat#: 113820, clone R17217 |
| CD45R (B220) – PerCP-Cyanine5.5 | Tonbo Biosciences | Cat#: 65-0452, clone RA3-6B2 |
| CD11c – BV510 | BioLegend | Cat#: 117353, clone N488 |
| CD4 – PE594 | BioLegend | Cat#: 100456, clone GK1.5 |
| CD8a – BV510 | BioLegend | Cat#: 100752, clone 53-6.7 |
| CD62L (L-Selectin) – PerCP-Cyanine 5.5 | Tonbo Biosciences | Cat#: 65-0621, clone MEL-14 |
| CD44 – RedFluor 710 | Tonbo Biosciences | Cat#: 80-0441, clone M7 |
| CD69 – PE-Cyanine7 | Tonbo Biosciences | Cat#: 60-0691, clone H1.2F3 |
| CD103 – APC | BioLegend | Cat#: 121414, clone 2E7 |
| CD3 - PE | BioLegend | Cat#: 100206, clone 17A2 |
| CD11b (Mac1) – FITC | Tonbo Biosciences | Cat#: 35-0112, clone M1/70 |
| CD3 – APC | Tonbo Biosciences | Cat#: 20-0032, clone 17A2 |
| CD4 – FITC | Tonbo Biosciences | Cat#: 35-0041, clone GK1.5 |
| CD25 – BV421 | BioLegend | Cat#: 102034, clone PC61 |
| CD11b (Mac1) – PE | Tonbo Biosciences | Cat#: 50-0112, clone M1/70 |
| CD45R (B220) – PE | BioLegend | Cat#: 103208, clone RA.3-6B2 |
| Ter119 – PE | BioLegend | Cat#: 116208, clone TER-119 |
| CD45.1 – VF450 | Tonbo Biosciences | Cat#: 75-0453, clone A20 |
| CD45.2 – PE/Cyanine7 | BioLegend | Cat#: 109830, clone 104 |
| CD45.2 – PE/Cyanine7 | Tonbo Biosciences | Cat#: 60-0454, clone 104 |
| Ly-6A/E (Sca-1) – PerCP/Cyanine5.5 | BioLegend | Cat#: 108124, clone D7 |
| CD2 – APC | BioLegend | Cat#: 100112, clone RM2.5 |
| CD5 – APC | BioLegend | Cat#: 100626, clone 53-7.3 |
| CD8a – APC | Tonbo Biosciences | Cat#: 20-0081, clone 53-6.7 |
| Ter119 – APC | Tonbo Biosciences | Cat#: 20-5921, clone TER-119 |
| CD45R (B220) – APC | Tonbo Biosciences | Cat#: 20-0452, clone RA.3-6B2 |
| Ly-6G (Gr-1) – APC | Tonbo Biosciences | Cat#: 20-5931, clone RB6-8C5 |
| CD34 – FITC | Invitrogen | Cat#: 11-0341-85, clone RAM34 |
| SiglecF – PE-CF594 | BD Biosciences | Cat#: 562757, clone E50-2440 |
| CD11b (Mac1) – BV510 | BioLegend | Cat#: 107263, clone M1/70 |
| CD45.1 – APC-Cyanine7 | Tonbo Biosciences | Cat#: 25-0453, clone A20 |

## Notes

### Competing Interest Statement

The authors have declared no competing interest.

## REFERENCES

1. Colley DG, Bustinduy AL, Secor WE, King CH. Human schistosomiasis. Lancet. 2014;383(9936):2253–64.

2. McManus DP, Dunne DW, Sacko M, Utzinger J, Vennervald BJ, Zhou XN. Schistosomiasis. Nat Rev Dis Primers. 2018;4(1):13.

3. Abdel Aziz N, Musaigwa F, Mosala P, Berkiks I, Brombacher F. Type 2 immunity: a two-edged sword in schistosomiasis immunopathology. Trends Immunol. 2022;43(8):657–73.

4. King CH. Parasites and poverty: the case of schistosomiasis. Acta Trop. 2010;113(2):95–104.

5. King CH, Dickman K, Tisch DJ. Reassessment of the cost of chronic helmintic infection: a meta-analysis of disability-related outcomes in endemic schistosomiasis. Lancet. 2005;365(9470):1561–9.

6. King CH, Dangerfield-Cha M. The unacknowledged impact of chronic schistosomiasis. Chronic Illn. 2008;4(1):65–79.

7. Lo NC, Addiss DG, Hotez PJ, King CH, Stothard JR, Evans DS, et al. A call to strengthen the global strategy against schistosomiasis and soil-transmitted helminthiasis: the time is now. Lancet Infect Dis. 2017;17(2):e64–e9.

8. Harris AR, Russell RJ, Charters AD. A review of schistosomiasis in immigrants in Western Australia, demonstrating the unusual longevity of Schistosoma mansoni. Trans R Soc Trop Med Hyg. 1984;78(3):385–8.

9. Fairfax K, Nascimento M, Huang SC, Everts B, Pearce EJ. Th2 responses in schistosomiasis. Semin Immunopathol. 2012;34(6):863–71.

10. Pearce EJ, MacDonald AS. The immunobiology of schistosomiasis. Nat Rev Immunol. 2002;2(7):499–511.

11. Girgis NM, Gundra UM, Loke P. Immune regulation during helminth infections. PLoS Pathog. 2013;9(4):e1003250.

12. Flammer PG, Ryan H, Preston SG, Warren S, Prichystalova R, Weiss R, et al. Epidemiological insights from a large-scale investigation of intestinal helminths in Medieval Europe. PLoS Negl Trop Dis. 2020;14(8):e0008600.

13. Shehata MA, Chama MF, Funjika E. Prevalence and intensity of Schistosoma haematobium infection among schoolchildren in central Zambia before and after mass treatment with a single dose of praziquantel. Trop Parasitol. 2018;8(1):12–7.

14. Toft II JD. The Pathoparasitology of Nonhuman Primates: A Review. In: Benirschke K, editor. Primates Proceedings in Life Sciences New York, NY: Springer; 1986.

15. Standley CJ, Mugisha L, Dobson AP, Stothard JR. Zoonotic schistosomiasis in non-human primates: past, present and future activities at the human-wildlife interface in Africa. J Helminthol. 2012;86(2):131–40.

16. Caiado F, Pietras EM, Manz MG. Inflammation as a regulator of hematopoietic stem cell function in disease, aging, and clonal selection. J Exp Med. 2021;218(7).

17. Matatall KA, Jeong M, Chen S, Sun D, Chen F, Mo Q, et al. Chronic Infection Depletes Hematopoietic Stem Cells through Stress-Induced Terminal Differentiation. Cell Rep. 2016;17(10):2584–95.

18. Baldridge MT, King KY, Boles NC, Weksberg DC, Goodell MA. Quiescent haematopoietic stem cells are activated by IFN-gamma in response to chronic infection. Nature. 2010;465(7299):793–7.

19. Burberry A, Zeng MY, Ding L, Wicks I, Inohara N, Morrison SJ, et al. Infection mobilizes hematopoietic stem cells through cooperative NOD-like receptor and Toll-like receptor signaling. Cell Host Microbe. 2014;15(6):779–91.

20. Morales-Mantilla DE, Kain B, Le D, Flores AR, Paust S, King KY. Hematopoietic stem and progenitor cells improve survival from sepsis by boosting immunomodulatory cells. Elife. 2022;11.

21. Haltalli MLR, Watcham S, Wilson NK, Eilers K, Lipien A, Ang H, et al. Manipulating niche composition limits damage to haematopoietic stem cells during Plasmodium infection. Nat Cell Biol. 2020;22(12):1399–410.

22. MacNamara KC, Jones M, Martin O, Winslow GM. Transient activation of hematopoietic stem and progenitor cells by IFNgamma during acute bacterial infection. PLoS One. 2011;6(12):e28669.

23. Abidin BM, Hammami A, Stager S, Heinonen KM. Infection-adapted emergency hematopoiesis promotes visceral leishmaniasis. PLoS Pathog. 2017;13(8):e1006422.

24. Yanez A, Murciano C, O’Connor JE, Gozalbo D, Gil ML. Candida albicans triggers proliferation and differentiation of hematopoietic stem and progenitor cells by a MyD88-dependent signaling. Microbes Infect. 2009;11(4):531–5.

25. Mistry JJ, Hellmich C, Moore JA, Jibril A, Macaulay I, Moreno-Gonzalez M, et al. Free fatty-acid transport via CD36 drives beta-oxidation-mediated hematopoietic stem cell response to infection. Nat Commun. 2021;12(1):7130.

26. Takizawa H, Fritsch K, Kovtonyuk LV, Saito Y, Yakkala C, Jacobs K, et al. Pathogen-Induced TLR4-TRIF Innate Immune Signaling in Hematopoietic Stem Cells Promotes Proliferation but Reduces Competitive Fitness. Cell Stem Cell. 2017;21(2):225–40 e5.

27. Isringhausen S, Mun Y, Kovtonyuk L, Krautler NJ, Suessbier U, Gomariz A, et al. Chronic viral infections persistently alter marrow stroma and impair hematopoietic stem cell fitness. J Exp Med. 2021;218(12).

28. Hirche C, Frenz T, Haas SF, Doring M, Borst K, Tegtmeyer PK, et al. Systemic Virus Infections Differentially Modulate Cell Cycle State and Functionality of Long-Term Hematopoietic Stem Cells In Vivo. Cell Rep. 2017;19(11):2345–56.

29. de Bruin AM, Demirel O, Hooibrink B, Brandts CH, Nolte MA. Interferon-gamma impairs proliferation of hematopoietic stem cells in mice. Blood. 2013;121(18):3578–85.

30. Inclan-Rico JM, Hernandez CM, Henry EK, Federman HG, Sy CB, Ponessa JJ, et al. Trichinella spiralis-induced mastocytosis and erythropoiesis are simultaneously supported by a bipotent mast cell/erythrocyte precursor cell. PLoS Pathog. 2020;16(5):e1008579.

31. Rashidi NM, Scott MK, Scherf N, Krinner A, Kalchschmidt JS, Gounaris K, et al. In vivo time-lapse imaging shows diverse niche engagement by quiescent and naturally activated hematopoietic stem cells. Blood. 2014;124(1):79–83.

32. Cortes-Selva D, Gibbs L, Maschek JA, Nascimento M, Van Ry T, Cox JE, et al. Metabolic reprogramming of the myeloid lineage by Schistosoma mansoni infection persists independently of antigen exposure. PLoS Pathog. 2021;17(1):e1009198.

33. Musaigwa F, Kamdem SD, Mpotje T, Mosala P, Abdel Aziz N, Herbert DR, et al. Schistosoma mansoni infection induces plasmablast and plasma cell death in the bone marrow and accelerates the decline of host vaccine responses. PLoS Pathog. 2022;18(2):e1010327.

34. Nunes-Alves C, Nobrega C, Behar SM, Correia-Neves M. Tolerance has its limits: how the thymus copes with infection. Trends Immunol. 2013;34(10):502–10.

35. Perlstein TS, Weuve J, Pfeffer MA, Beckman JA. Red blood cell distribution width and mortality risk in a community-based prospective cohort. Arch Intern Med. 2009;169(6):588–94.

36. Friedman JF, Kanzaria HK, McGarvey ST. Human schistosomiasis and anemia: the relationship and potential mechanisms. Trends Parasitol. 2005;21(8):386–92.

37. Butler SE, Muok EM, Montgomery SP, Odhiambo K, Mwinzi PM, Secor WE, et al. Mechanism of anemia in Schistosoma mansoni-infected school children in Western Kenya. Am J Trop Med Hyg. 2012;87(5):862–7.

38. Ganz T. Anemia of Inflammation. N Engl J Med. 2019;381(12):1148–57.

39. Leite LA, Pimenta Filho AA, Ferreira Rde C, da Fonseca CS, dos Santos BS, Montenegro SM, et al. Splenectomy Improves Hemostatic and Liver Functions in Hepatosplenic Schistosomiasis Mansoni. PLoS One. 2015;10(8):e0135370.

40. Tamarozzi F, Fittipaldo VA, Orth HM, Richter J, Buonfrate D, Riccardi N, et al. Diagnosis and clinical management of hepatosplenic schistosomiasis: A scoping review of the literature. PLoS Negl Trop Dis. 2021;15(3):e0009191.

41. Bach O, Baier M, Pullwitt A, Fosiko N, Chagaluka G, Kalima M, et al. Falciparum malaria after splenectomy: a prospective controlled study of 33 previously splenectomized Malawian adults. Trans R Soc Trop Med Hyg. 2005;99(11):861–7.

42. Rossi MI, Dutra HS, El-Cheikh MC, Bonomo A, Borojevic R. Extramedullar B lymphopoiesis in liver schistosomal granulomas: presence of the early stages and inhibition of the full B cell differentiation. Int Immunol. 1999;11(4):509–18.

43. Dutra HS, Rossi MI, Azevedo SP, el-Cheikh MC, Borojevic R. Haematopoietic capacity of colony-forming cells mobilized in hepatic inflammatory reactions as compared to that of normal bone marrow cells. Res Immunol. 1997;148(7):437–44.

44. el-Cheikh MC, Borojevic R. Extramedullar proliferation of eosinophil granulocytes in chronic schistosomiasis mansoni is mediated by a factor secreted by inflammatory macrophages. Infect Immun. 1990;58(3):816–21.

45. Maruyama H, Higa A, Asami M, Owhashi M, Nawa Y. Extramedullary eosinophilopoiesis in the liver of Schistosoma japonicum-infected mice, with reference to hemopoietic stem cells. Parasitol Res. 1990;76(6):461–5.

46. Borojevic R, Nicola MH, Santos-da-Silva C, Grimaldi G, Jr. Schistosoma mansoni: extramedullar eosinophil myelopoiesis induced by intraperitoneal glass implants in chronically infected mice. Exp Parasitol. 1985;59(3):290–9.

47. Borojevic R, Stocker S, Grimaud JA. Hepatic eosinophil granulocytopoiesis in murine experimental Schistosomiasis mansoni. Br J Exp Pathol. 1981;62(5):480–9.

48. Francisco JS, Terra M, Klein GCT, Dias de Oliveira B, Pelajo-Machado M. The hepatic extramedullary hematopoiesis during experimental murine Schistosomiasis mansoni. Front Immunol. 2022;13:955034.

49. Gundra UM, Girgis NM, Gonzalez MA, San Tang M, Van Der Zande HJP, Lin JD, et al. Vitamin A mediates conversion of monocyte-derived macrophages into tissue-resident macrophages during alternative activation. Nat Immunol. 2017;18(6):642–53.

50. Nascimento M, Huang SC, Smith A, Everts B, Lam W, Bassity E, et al. Ly6Chi monocyte recruitment is responsible for Th2 associated host-protective macrophage accumulation in liver inflammation due to schistosomiasis. PLoS Pathog. 2014;10(8):e1004282.

51. Girgis NM, Gundra UM, Ward LN, Cabrera M, Frevert U, Loke P. Ly6C(high) monocytes become alternatively activated macrophages in schistosome granulomas with help from CD4+ cells. PLoS Pathog. 2014;10(6):e1004080.

52. Brito JM, Borojevic R. Liver granulomas in schistosomiasis: mast cell-dependent induction of SCF expression in hepatic stellate cells is mediated by TNF-alpha. J Leukoc Biol. 1997;62(3):389–96.

53. Correia AL, Guimaraes JC, Auf der Maur P, De Silva D, Trefny MP, Okamoto R, et al. Hepatic stellate cells suppress NK cell-sustained breast cancer dormancy. Nature. 2021;594(7864):566–71.

54. Lee Y, Leslie J, Yang Y, Ding L. Hepatic stellate and endothelial cells maintain hematopoietic stem cells in the developing liver. J Exp Med. 2021;218(3).

55. Tucker MS, Karunaratne LB, Lewis FA, Freitas TC, Liang YS. Schistosomiasis. Curr Protoc Immunol. 2013;103:19 1 1–1 58.

